# Differences in intrinsic cellular O-GlcNAcylation impact response to metabolic stress

**DOI:** 10.1101/2025.10.24.684323

**Authors:** Sadia Rahmani, Melissa Huestis, Warren W. Wakarchuk, Costin N. Antonescu

## Abstract

The cellular response to metabolic stress is complex and involves various signals and pathways leading to cellular adaptation. AMP-activated Protein Kinase (AMPK) senses nutrient insufficiency reflected in the ratio of ATP to AMP/ADP and/or glucose availability. In addition, the dynamic modification of proteins with O-linked β-N-Acetylglucosamine (O-GlcNAc) by O-GlcNAc transferase (OGT) is also nutrient sensitive, given that the substrate for this modification is the product of the hexosamine biosynthetic pathway. AMPK can also regulate the hexosamine biosynthetic pathway, and O-GlcNAc may reciprocally regulate AMPK activity under some contexts. However, it remains unclear which parameters establish the extent of reciprocal regulation of AMPK and protein O-GlcNAc modification and whether these signals can be orthogonal under some circumstances. Here, we used ARPE-19 and MDA-MB-231 cells to examine how nutrient limitation or pharmacological activation combined with inhibition of AMPK and O-GlcNAcylation impacts the activation of the reciprocal pathway. We found that the intrinsic level of global protein O-GlcNAc modification may impact how AMPK and metabolic stress regulate global O-GlcNAcylation, and that both signaling systems could be orthogonal in some contexts. These results contribute to understanding the complexity of cellular signaling in response to nutrient availability.

## Introduction

Various cells and tissues routinely experience some form of intermittent nutrient limitation that requires time-sensitive adaptation to ensure appropriate cellular and tissue function (Fulda et al. 2010; Martinez et al. 2017). Nutrient limitation triggers changes to specific signaling pathways that include AMP-Activated Kinase (AMPK) and O-linked ß-N-acetylglucosamine (O-GlcNAc) transferase (OGT), which changes the activity and function of many different proteins to allow this metabolic adaptation. While AMPK and OGT control at least partly unique subsets of proteins, both of these essential regulatory enzymes respond to similar cues (e.g. availability of glucose and other nutrients) (Yao et al. 2018), suggesting the existence of signaling cross-talk to facilitate cellular adaptation to nutrient conditions (Bond and Hanover 2015). Specifically, reciprocal regulation of global O-GlcNAc modification and AMPK activity has been proposed to allow a cell to coordinate the action of these signals during cellular adaptation to nutrient limitation (Cork et al. 2018). The extent to which AMPK and OGT signals undergo reciprocal regulation upon nutrient stress remains incompletely understood.

AMPK is a heterotrimeric protein composed of two regulatory (ß, γ) and one catalytic (α) subunits which functions as a serine/threonine kinase (Lin and Hardie 2018). AMPK activity in nutrient-replete conditions is low, and this kinase becomes activated upon a reduction in intracellular ATP:ADP (or ATP:AMP) ratio sensed by direct binding of nucleoside phosphates to the AMPK y subunit (Yan et al. 2018) or a reduction in glycolytic flux which is sensed indirectly by levels of fructose-1,6-bisphosphate (Zhang et al. 2017). AMPK activation by these nutrient sensing mechanisms leads to enhanced phosphorylation at amino acid T172 within the a subunit (Willows et al. 2017). In response to metabolic stress, AMPK controls many cellular processes by phosphorylation of specific substrates, including the downregulation of several ATP-consuming anabolic reactions and upregulation of nutrient uptake and some catabolic reactions that enhance generation of ATP. For example, AMPK phosphorylation of acetyl-CoA carboxylase (ACC) impairs fatty acid synthesis and enhances fatty acid fl-oxidation (Lee et al. 2018).

The OGT enzyme catalyzes the modification of Ser/Thr residue of a wide range of proteins with an uncharged bulky O-linked ß-N-acetylglucosamine (Lazarus et al. 2011), a modification that can be removed by action of a specific O-GlcNAc hydrolase (OGA) (Roth et al. 2017). In contrast to complex glycans on the exofacial portion of cell surface proteins, O-GlcNAc modification of proteins within the cytosol controlled by OGT and OGA is highly dynamic and regulated by a number of different signals and cues (Bond and Hanover 2015). O-GlcNAcylation impacts the function of target proteins in many different ways specific to the protein being modified, which includes changes to protein half-life, enzyme activity, subcellular localization and various protein-protein interactions (Tarbet et al. 2018). While some targeting of specific substrates occurs, OGT impacts a wide range of substrates through this modification (Sprung et al. 2005; Alfaro et al. 2012). This broad action of OGT on a wide variety of protein substrates is in contrast to the activity of specific kinases such as AMPK, which modify a restricted set of substrates with defined consensus sequences (Ducommun et al. 2015). As such, global (total cellular) O-GlcNAcylation of protein is an effective readout of OGT/OGA activity in a cell (Zachara et al. 2004; Swamy et al. 2016; Zhu et al. 2018; Gélinas et al. 2018).

The donor substrate for protein O-GlcNAcylation is uridine diphosphate N-acetylglucosamine (UDP-GlcNAc). Importantly, UDP-GlcNAc is the product of the hexosamine biosynthetic pathway that is in turn controlled by the input of glucose, glutamine and other metabolites (Abdel Rahman et al. 2013). The level of UDP-GlcNAc can be limiting for O-GlcNAc modification of proteins (Kreppel and Hart 1999), resulting in control of the global O-GlcNAcylation of proteins by the hexosamine biosynthetic pathway. As a result, global O-GlcNAc protein modification acts as a key nutrient-sensor for fluctuations in glucose, glutamine and other metabolites (Bond and Hanover 2015).

Based solely on how nutrient availability impacts UDP-GlcNAc levels and thus O-GlcNAcylation by OGT, as well as how nutrient availability controls AMPK activation, an inverse correlation between the extent of these two post-translational modifications (PTMs) would be expected. For example, considering only the direct regulation of each of global protein O-GlcNAcylation and AMPK activity under the metabolic stress of glucose limitation, AMPK activity should be enhanced, while broad protein O-GlcNAcylation should be decreased, whereas under conditions of metabolic sufficiency, AMPK activity should be suppressed, and broad protein O-GlcNAcylation should be enhanced. However, several lines of evidence suggest that broad protein O-GlcNAcylation and AMPK-dependent protein phosphorylation exhibit signaling cross-talk and reciprocal regulation during acute (<4h) nutrient limitation. While additional reciprocal regulation of global protein O-GlcNAcylation and AMPK can occur upon chronic nutrient limitation, for example involving changes in expression of AMPK and/or OGT (Cheung and Hart 2008), we focus here on the signaling elicited by acute (<4h) metabolic stress.

Several studies indicate that AMPK can acutely regulate global O-GlcNAc modification via direct phosphorylation of enzymes critical for dynamic O-GlcNAcylation. AMPK directly phosphorylates glutamine-fructose-6-phosphate transaminase 1 (GFAT1) at Ser243, resulting in suppression of the hexosamine biosynthetic pathway upon AMPK activation during metabolic stress (Eguchi et al. 2009; Zibrova et al. 2017; Gélinas et al. 2018). In contrast, several studies have instead reported that many different cellular stressors, including metabolic stressors that also activate AMPK activation trigger enhancement of global protein O-GlcNAcylation (Martinez et al. 2017). This suggests that additional mechanisms exist that trigger enhancement of global protein O-GlcNAcylation as a result of, or in parallel to, AMPK activation. Hence, AMPK and/or other metabolic stress signals might elicit enhancement or suppression of global protein O-GlcNAcylation under different contexts. The molecular and cellular parameters that define how AMPK controls global protein O-GlcNAcylation remain poorly understood.

In addition to the evidence for context-specific regulation of global O-GlcNAc modification by AMPK and/or other metabolic stress signals, OGT can O-GlcNAc modify AMPK on the α1-, α2-, γ1-, γ2-, and γ3-subunits, resulting in control of AMPK activity (Bullen et al. 2014). Blunting of OGA activity by acute treatment (<4 h) with the OGA inhibitor Thiamet G results in reduced AMPK Thr172 phosphorylation and reduced AMPK-dependent ACC phosphorylation in C_2_C_12_ myotubes (Bullen et al. 2014). The regulation of AMPK by O-GlcNAc appears to be complex, as O-GlcNAcylation of AMPK requires AMPK activity. Consistent with this, silencing of OGT in MDA-MB-231 cells enhances AMPK phosphorylation (Sodi et al. 2018) and overexpression of OGT in LoVo cells suppresses AMPK and ACC phosphorylation (Ishimura et al. 2017). In contrast to these studies indicating that OGT and O-GlcNAc modification can regulate AMPK, other studies have found no effect of alterations of global O-GlcNAc modification on AMPK. Perfusion of isolated rat hearts with glucosamine enhanced UDP-GlcNAc levels and global protein O-GlcNAcylation but did not impact AMPK or ACC phosphorylation (Laczy et al. 2011), and enhancement or suppression of UDP-GlcNAc levels had no effect on AMPK activity in rat cardiomyocytes or rat hearts (Gélinas et al. 2018). Thus, the regulation of AMPK by O-GlcNAc modification may also be context-specific.

From these studies emerges the notion that there may be cell context-specific parameters that define the extent and nature of the reciprocal control of global protein O-GlcNAcylation and AMPK activity. Importantly, several factors can lead to distinct levels of global protein O-GlcNAcylation in different cellular contexts. For example, during T cell development and differentiation, the enhancement of glucose and glutamine uptake at specific stages of development leads to enhanced OGT-mediated global protein O-GlcNAcylation (Swamy et al. 2016). This indicates that the relative contribution of each of glucose and glutamine uptake, channelling of metabolism into the hexosamine biosynthetic pathway (HBP) as determined by GFAT1 expression and/or activity, and expression of OGT and OGA may shift the limiting factor controlling global O-GlcNAcylation among different cell lines. As a result, cells that exhibit intrinsically high global protein O-GlcNAcylation that nears cellular saturation for this modification may exhibit distinct reciprocal regulation of global protein O-GlcNAcylation and AMPK activation than cells with lower intrinsic global protein O-GlcNAcylation. Resolving the extent of direct reciprocal signaling regulation of AMPK and global protein O-GlcNAc modification in specific contexts requires approaches that can observe the effects of acute (<4h) perturbation or activation of AMPK, OGT and OGA.

We thus hypothesize that the extent of the intrinsic global O-GlcNAcylation in a specific cell type establishes the extent to which global O-GlcNAcylation and AMPK exhibit reciprocal regulation, making this signaling cross-talk possible but not obligatory in all cells. To examine this, we first compare the ability of acute (<4h) inhibition of OGT and OGA in two different cell lines, ARPE-19 (a non-immortalized human epithelial cell line) and MDA-MB-231 (a triple negative breast cancer cell line). We find that these cell lines differ in the extent of initial global O-GlcNAcylation observed and in the extent to which OGA inhibition impacts global protein O-GlcNAcylation. We then find that ARPE-19 cells, which exhibit lower intrinsic global O-GlcNAcylation, exhibit substantial enhancement of O-GlcNAcylation upon acute OGA inhibition as well as a significant increase of global protein O-GlcNAcylation in response to metabolic stress signals. In contrast, MDA-MB-231 cells exhibit higher intrinsic global protein O-GlcNAcylation that is modestly responsive to OGA inhibition. Importantly, MDA-MB-231 cells exhibit no change in O-GlcNAcylation in response to metabolic stress signals. We also observe that inhibition of OGT or OGA do not impact AMPK activation upon metabolic stress, suggesting that in some contexts, AMPK and global protein O-GlcNAcylation can be orthogonal metabolic signaling pathways.

## MATERIALS AND METHODS

### Materials

Details about the materials used in this study can be found in the Supporting Information (Table S1).

### Cell Culture

Wild-type human retinal pigment epithelial (ARPE-19) and triple negative breast cancer (MDA-MB-231) cells were cultured as previously described (Bautista et al. 2018) in Dulbecco’s Modified Eagle’s Medium/F-12 (DMEM/F12; Thermo Fisher Scientific, Mississauga, ON) supplemented with 10% fetal bovine serum (Thermo Fisher Scientific), and an antibiotic cocktail consisting of 100 U/mL penicillin and 100 μg/mg streptomycin (Thermo Fisher Scientific). Cells were incubated at 37C and 5% CO_2_ and 95% O_2_ until they reached 80% confluency and were suitable for passaging. Cells were passaged every 2-3 days, washed with sterile phosphate buffered saline (Sigma Aldrich, Oakville, ON) and lifted with 0.25% Trypsin-EDTA (Thermo Fisher Scientific).

### Pharmacological treatments

Prior to all experiments, ARPE-19 and MDA-MB-231 cells were washed and incubated in low serum media (DMEM/F12 supplemented with 0.1% fetal bovine serum) for 1h prior to any incubation with OGT/OGA inhibitors or metabolic stress treatments. Details of OGT/OGA inhibitors and metabolic stress treatments used in each experiment can be found in the corresponding figure legend.

### Western blotting

Western blotting was performed as previously described (Garay et al. 2015; Delos Santos et al. 2017). After incubation and pharmacological treatments as indicated, cells were rapidly cooled by washing with ice-cold PBS and then lysed in in 2X-Laemelli Sample buffer (0.5M Tris pH 6.8, Glycerol, 10% SDS) supplemented with 1 mM sodium orthovanadate, 10 mM okadaic acid and 20 mM of protease inhibitor. Whole-cell lysates were then syringed 5 times with a 27.5-gauge syringe. Lysates were then heated at 65°C for 15 minutes under reducing conditions (supplementation of lysis buffer with 10% beta-mercaptoethanol and 5% bromophenol blue), resolved by SDS-PAGE, and transferred to 0.2 m pore PVDF membrane (PALL Life Science, Port Washington). The membrane was blocked with 3% BSA and then incubated with indicated antibodies in 1% BSA at 1:1000 dilution at 4C overnight. The list of primary and secondary antibodies is in Table S1. The membrane was then subjected to a secondary HRP-conjugated antibody (with 1% BSA) and left to shake for 1 hour at RT before imaging. Bands were visualized with Luminata ECL chemiluminescence substrate (Milipore Sigma, Oakville, ON) and quantified by densitometry analysis with ImageJ (version 1.50, National Institutes of Health, Bethesda, MD) (Schneider et al. 2012) to quantify protein abundance through the detection of differences in band signal intensities. Quantification of global protein O-GlcNAcylation was done by densitometry of the entire lane in each condition.

### Statistical analysis

All results are represented as mean ± standard error of the mean (SEM). Statistical analysis was performed with Prism 7 for Mac OS X (version 7.2, GraphPad Software, Inc.) and is described in detail in each figure caption. Values for p < 0.05 were considered statistically significant.

## RESULTS

### MDA-MB-231 cells may have intrinsic global protein O-GlcNAcylation closer to saturation than ARPE-19 cells

To determine if different cell lines exhibit differences in the dynamic regulation of global protein O-GlcNAcylation, we examined MDA-MB-231 and ARPE-19 cells. We measured whole-cell O-GlcNAcylation by immunoblotting whole-cell lysates using the RL2 O-GlcNAc antibody that recognizes an epitope containing (serine/threonine) O-linked ß-N-acetylglucosamine and can thus recognize a wide range of proteins when O-GlcNAc modified. Western blotting of whole-cell lysates reveals the typical large range of proteins since the O-GlcNAc modification can occur on a number of intracellular proteins (**Fig. S1**).

To determine how global O-GlcNAcylation is impacted by acute suppression of either OGT or OGA, we treated each cell line with a OGT inhibitor (peracetylated 2-acetamido-2-deoxy-**5**-**thio**-D-glucopyranose, Ac-5SGlcNAc) (Gloster et al. 2011) or with an OGA inhibitor (Thiamet G) (Yuzwa et al. 2008) for various times up to 4 h (**Fig. 1**). OGT inhibition by 50 µM Ac-5SGlcNAc treatment resulted in a similar time-dependent reduction of whole-cell O-GlcNAc modification in both cell lines, with a ∼50% decrease in whole-cell O-GlcNAcylation by 4 h of treatment with this inhibitor (**Fig. 1C**).

**Figure 1.**
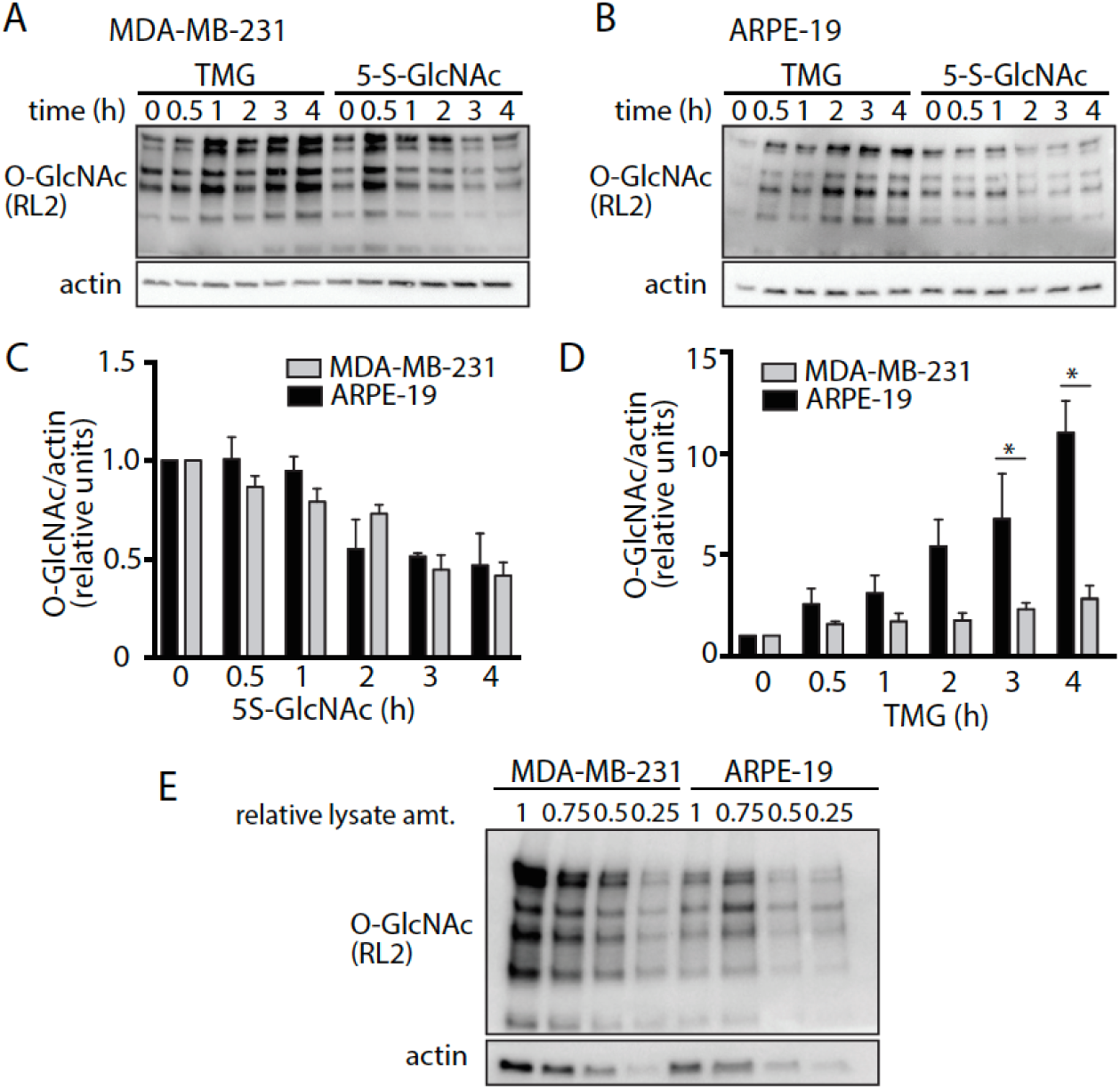
Distinct impact of OGA inhibition in MDA-MB-231 and ARPE-19 cells. MDA-MB-231 cells or ARPE-19 cells were incubated in low serum media for 1 h before being stimulated with either 50 µM 5S-GlcNAc (OGT inhibitor) or 20 µM Thiamet G (TMG, OGA inhibitor) for 0-4 h, as indicated. (**A**) MDA-MB-231 or (**B**) ARPE-19 whole-cell lysates were immunoblotted for O-GlcNAc modified proteins (RL2 antibody) and actin (loading control); shown are representative immunoblots. Shown are the mean means ± S.E. of the global protein O-GlcNAcylation detected in each cell type upon treatment with 5S-GlcNAc (**C**) or Thiamet G (**D**). Values are means ± S.E. *n =* 3 for all data sets quantified. *, p<0.05, determined by two-way ANOVA and Sidak’s multiple comparisons post-hoc test. (**E**) Shown are representative immunoblots of a range of different relative amounts of whole cell lysates from basal (non-drug-treated) MDA-MB-231 or ARPE-19 cells, detecting O-GlcNAc modified proteins (RL2 antibody) and actin (loading control).

Interestingly, and in contrast to the similar effect of OGT inhibition observed in both cell lines, OGA inhibition with 20 µM Thiamet G exhibited a dramatic difference in magnitude of response between cell lines. In ARPE-19 cells, Thiamet G treatment resulted in a dramatic time-dependent increase in whole-cell O-GlcNAcylation, with a 11.0 ± 2.2 fold increase by 4 h (n = 3, p < 0.01, **Fig. 1D**). In contrast, in MDA-MB-231 cells, Thiamet G treatment resulted in much more modest enhancement of whole-cell O-GlcNAcylation, resulting in only 2.8 ± 1.1 fold increase in this modification by 4h (n = 3, p < 0.01, **Fig. 1D**).

This difference in response to OGA inhibition between ARPE-19 and MDA-MB-231 cells could be due to several factors, including either 1) an intrinsically higher OGT activity in ARPE-19 cells, triggering an elevation in both initial and Thiamet G-induced gain in global protein O-GlcNAc modification or 2) a higher intrinsic level of global O-GlcNAcylation in MDA-MB-231 cells that may be closer to saturation than in ARPE-19 cells. To resolve this, we compared lysates from control (not treated with inhibitors) MDA-MB-231 and ARPE-19 cells, which revealed that MDA-MB-231 cells had elevated intrinsic global O-GlcNAc levels compared to ARPE-19 cells (**Fig. 1E**). This suggests that it is unlikely that ARPE-19 cells have intrinsically higher O-GlcNAc modification activity catalyzed by OGT than MDA-MB-231 cells, and instead indicate that MDA-MB-231 cells have a higher initial extent of global O-GlcNAc modification than ARPE-19 cells. As such, the difference in the increase in whole-cell O-GlcNAc modification upon treatment with Thiamet G may be due to MDA-MB-231 cells being closer to saturating global protein O-GlcNAcylation than ARPE-19 cells. Thus, MDA-MB-231 cells may be similar in this regard to T cells, which also have been described as having near-saturating levels of global protein O-GlcNAcylation (Swamy et al. 2016).

### AMPK-dependent changes in global protein O-GlcNAcylation may be defined by initial, intrinsic levels of O-GlcNAcylation

To determine if the initial level of global protein O-GlcNAcylation may impact the cellular response of whole cell O-GlcNAcylation to metabolic stressors that activate AMPK, we investigated the effect of treatment with the AMPK activator A769662 (A7) or 2-deoxyglucose (2DG, a non-metabolizable glucose analog that impairs glycolytic flux, which increases the intracellular AMP:ATP ratio) in ARPE-19 and MDA-MB-231 cells. Treatment of ARPE-19 cells with either 2DG or A7 triggered a significant increase in global protein O-GlcNAcylation (**Fig. 2A**). This suggests that ARPE-19 cells may have the capacity to enhance global O-GlcNAcylation, perhaps due to a relatively low initial extent of this modification.

**Figure 2.**
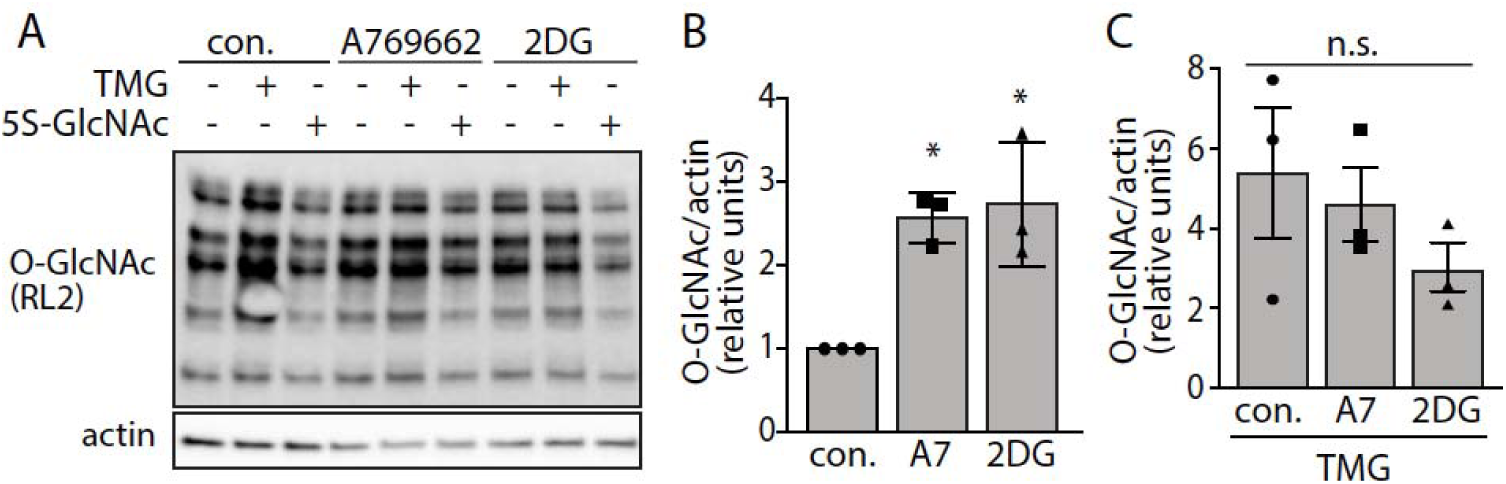
ARPE-19 cells increase whole cell O-GlcNAcylation upon treatment with metabolic stress stimuli. ARPE-19 cells were incubated in low serum and then treated with 20 µM Thiamet G (TMG) for 4 h or 50 µM Ac-5SGlcNAc (5S-GlcNAc) for 4 h, or left without treatment with OGA/OGT inhibitors (control, con.), after which cells were then stimulated with either 25 mM 2-deoxyglucose (2DG) for 20 min or 100 µM A769662 (A7) for 1 h in the continued presence of OGA/OGT inhibitors. (**A**) Shown are representative immunoblots detecting O-GlcNAc modified proteins (RL2 antibody) and actin (loading control). Also shown are the means ± S.E. (bars) as well as individual experiment measurements (*n =* 3) for (**B**) treatment with A7 or 2DG only (not treated Ac-5S-GlcNAc or Thiamet G) or (**C**) treatment with Thiamet G but alongside A7 or 2DG. *, p<0.05 determined by one-way ANOVA with Sidak’s multiple comparisons post-hoc test.

To determine the extent of the response of ARPE-19 cells to metabolic stressors upon an elevation of initial global O-GlcNAcylation, we compared the effect of A7 and 2DG treatment in cells that had been pre-treated with the OGA inhibitor Thiamet G for 4 h. This Thiamet G pre-treatment results in global protein O-GlcNAcylation in ARPE-19 cells that may more closely resemble the higher initial levels of O-GlcNAcylation in untreated MDA-MB-231 cells. Accordingly, treatment of ARPE-19 cells with 2DG or A7 in Thiamet G-pretreated cells resulted in no significant changes in global protein O-GlcNAcylation (**Fig. 2B**).

In contrast to the effects observed in ARPE-19 cells, treatment of MDA-MB-231 cells with either of these agents, or additional agents that activate AMPK, such as H_2_O_2_ or 5 µM oligomycin (OG) for 20 min resulted in no detectable difference in global protein O-GlcNAcylation (**Fig. 3**). While no changes in global protein O-GlcNAcylation were detected upon treatment with various metabolic stressors, we indeed observed strong activation of AMPK, detected by phosphorylation of the AMPK substrate acetyl-CoA carboxylase (ACC) (**Fig. 3**). These results indicate that one of the parameters that establishes the cell context-dependent regulation of global O-GlcNAcylation by metabolic signals that activate AMPK may be the initial, intrinsic levels of global protein O-GlcNAcylation.

**Figure 3.**
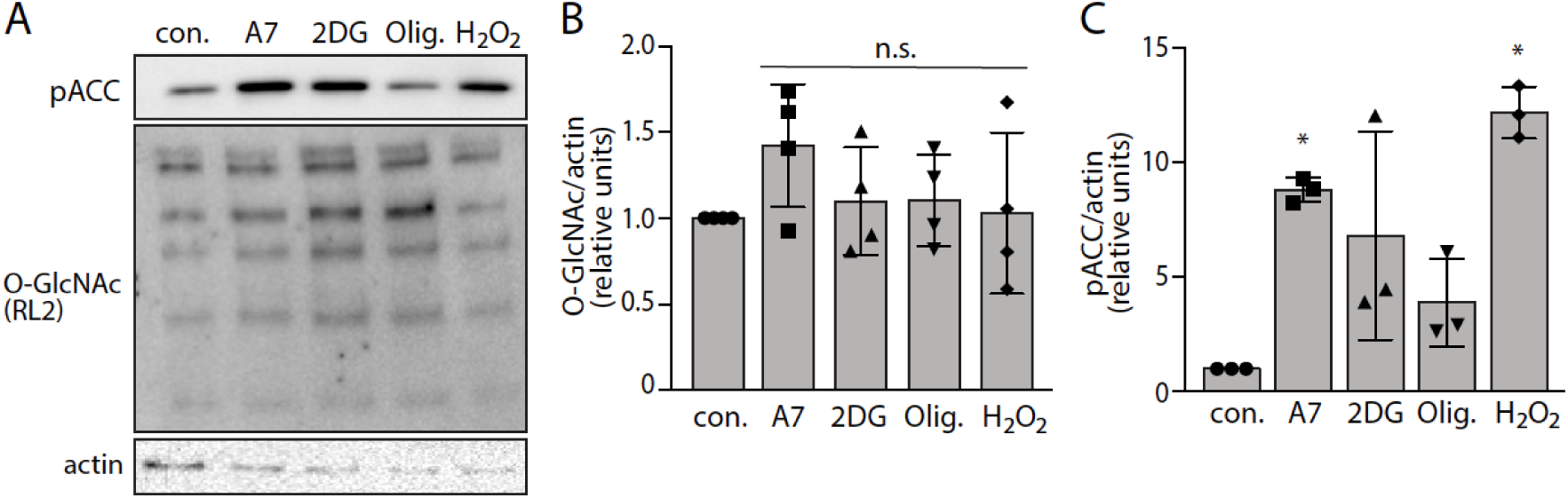
MDA-MB-231 cells did not change O-GlcNAcylation upon treatment with metabolic stress stimuli. MDA-MB-231 cells were incubated in low serum media and then stimulated with either 25 mM 2-deoxyglucose (2DG) for 20 min, 100 µM A769662 (A7) for 1 h, 5 µM oligomycin (Olig.) for 20 min, or 1 mM H_2_O_2_ for 20 min. (**A**) Shown are representative immunoblots detecting O-GlcNAc modified proteins (RL2 antibody) and phosphorylated ACC (pACC). Also shown are the means ± S.E. (bars) as well as individual experiment measurements for measurements of (**B**) total protein O-GlcNAcylation (*n* = 4) or (**C**) pACC levels. n.s. (not significant) (*n* = 3) and *, p<0.05 determined by one-way ANOVA with Sidak’s multiple comparisons post-hoc test.

### AMPK activation is independent of OGT and OGA activity in ARPE-19 and MBA-MB-231 cells

Next, we sought to determine if AMPK activation is altered upon stress stimuli in a matter that depends on cellular O-GlcNAcylation. To do so, we assessed the phosphorylation of ACC, a ubiquitous substrate of AMPK as a measure of AMPK activity. AMPK was robustly activated upon treatment with various metabolic stress signal and AMPK activation inducers in both ARPE-19 and MDA-MB-231 cells compared to the respective basal controls (**Fig. 4**). That these two cell lines differ in initial, intrinsic global protein O-GlcNAc levels yet exhibit comparable AMPK activation is consistent with the notion that AMPK activation may be independent of the extent of global cellular O-GlcNAcylation. To specifically probe how global O-GlcNAcylation impacts AMPK activation, we pre-treated cells with inhibitors of OGT or OGA for 4 hours (50 µM Ac-5SGlcNAc and 20 µM Thiamet G, respectively) before treatment with metabolic stress signals. Interestingly, inhibition of either OGT or OGA did not change the level of AMPK activation by a variety of stress stimuli (A7, 2DG, H_2_O_2_) (**Fig. 4**). These conditions of OGT and OGA inhibition (50 µM Ac-5SGlcNAc and 20 µM Thiamet G for 4h) indeed elicited robust changes in total cellular O-GlcNAcylation (**Fig. 1**), indicating that these inhibitors are effective at suppressing their respective enzyme targets. This indicates that unlike the suppression of AMPK activation by inhibition of OGA with Thiamet G (4-8h) in C_2_C_12_ myotubes (Bullen et al. 2014), suppression of either OGT or OGA for 4h has no effect on AMPK activity in ARPE-19 and MDA-MB-231 cells. This suggests that AMPK activation by acute metabolic stress signals (<1h) is not rigidly controlled by changes in global O-GlcNAcylation, and instead indicates that control of AMPK activation by O-GlcNAcylation may be restricted to specific cell types and/or longer durations (>4h) of perturbation of O-GlcNAcylation.

**Figure 4.**
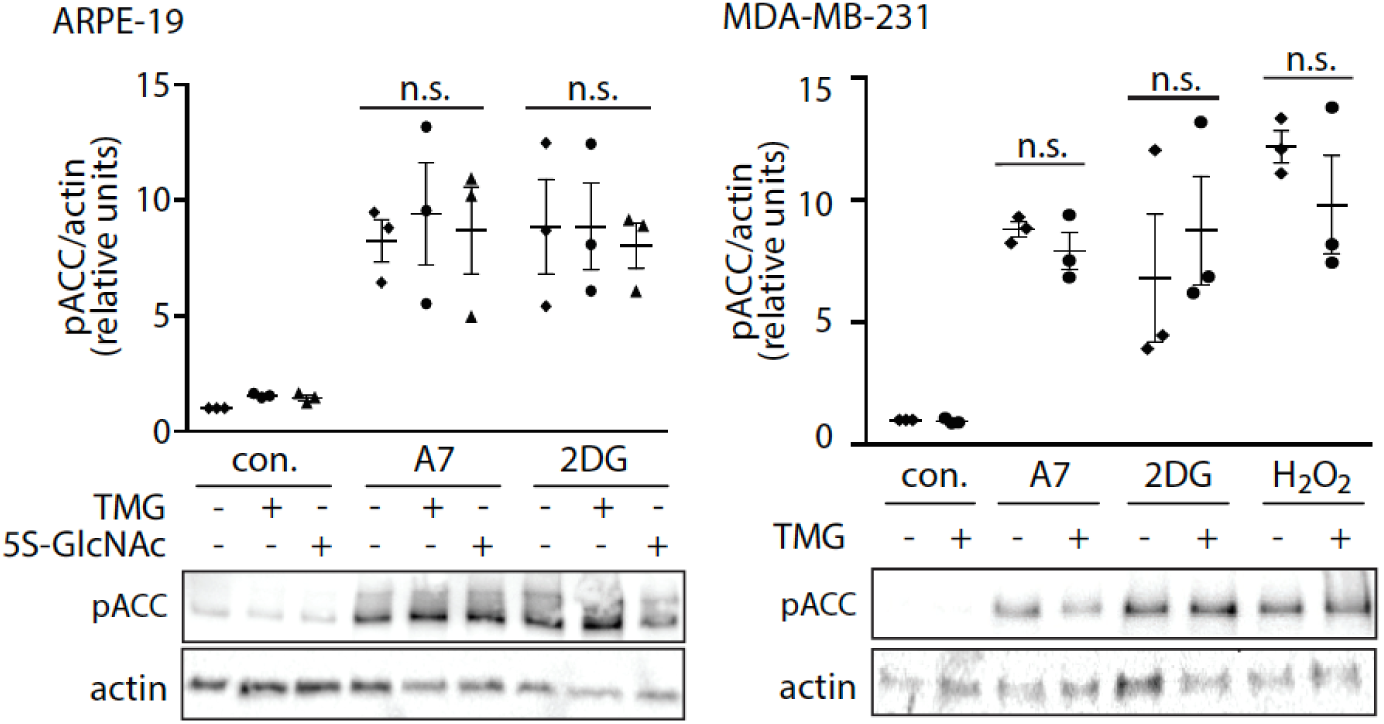
AMPK activation upon treatment with metabolic stress triggers is largely unaffected by inhibition of OGT or OGA. ARPE-19 cells or MDA-MB-231 cells were incubated in low serum media for 1 h before being treated with either 50 µM Ac-5SGlcNAc or 20 µM Thiamet G for 4 hrs. Cells were then stimulated with either 25 mM 2-deoxyglycose (2DG) for 20 min, 100 µM A769662 (A7) for 1 h, or 1 mM H_2_O_2_ for 20 min in the continued presence (or not) of OGA/OGT inhibitors. Shown (bottom panels) are representative immunoblots of phosphorylated ACC (pACC) or actin (loading control). Also shown are mean ± S.E. *n =* 3 (bars) as well as individual experiment measurements (*n =* 3) of pACC levels (top panels). n.s. (not significant), based on p<0.05 threshold, determined by two-way ANOVA and Sidak’s multiple comparisons post-hoc test.

### Metabolic-stress induced p38 MAPK activation is largely independent of OGT and OGA

In addition to the activation of AMPK, metabolic stress also leads to the activation of other stress signals. We next sought to determine if the extent of cellular global O-GlcNAcylation or the activity of OGT and OGA may regulate the activation of other stress signaling pathways that are activated in response to metabolic stress signals. To do so, we examined the requirement for phosphorylation of p38 mitogen activated protein kinase (p38 MAPK) or c-Jun NH_2_-terminal kinase (JNK). Treatment of MDA-MB-231 cells with specific stress stimuli such as A7, 2DG, oligomycin and H_2_O_2_ did not appreciably alter JNK phosphorylation, but each of these treatments did significantly increase p38 MAPK activation for all stress types (assessed by immunoblotting for phosphorylation of T180/Y182 on p38 MAPK). Pre-treatment of MDA-MB-231 cells with the OGA inhibitor Thiamet G did not change the activation of p38 MAPK by the different stressors (**Fig. 5A-C**). suggesting that like AMPK activation, p38 MAPK phosphorylation in response to metabolic stressors is not altered by enhancement of O-GlcNAcylation.

**Figure 5.**
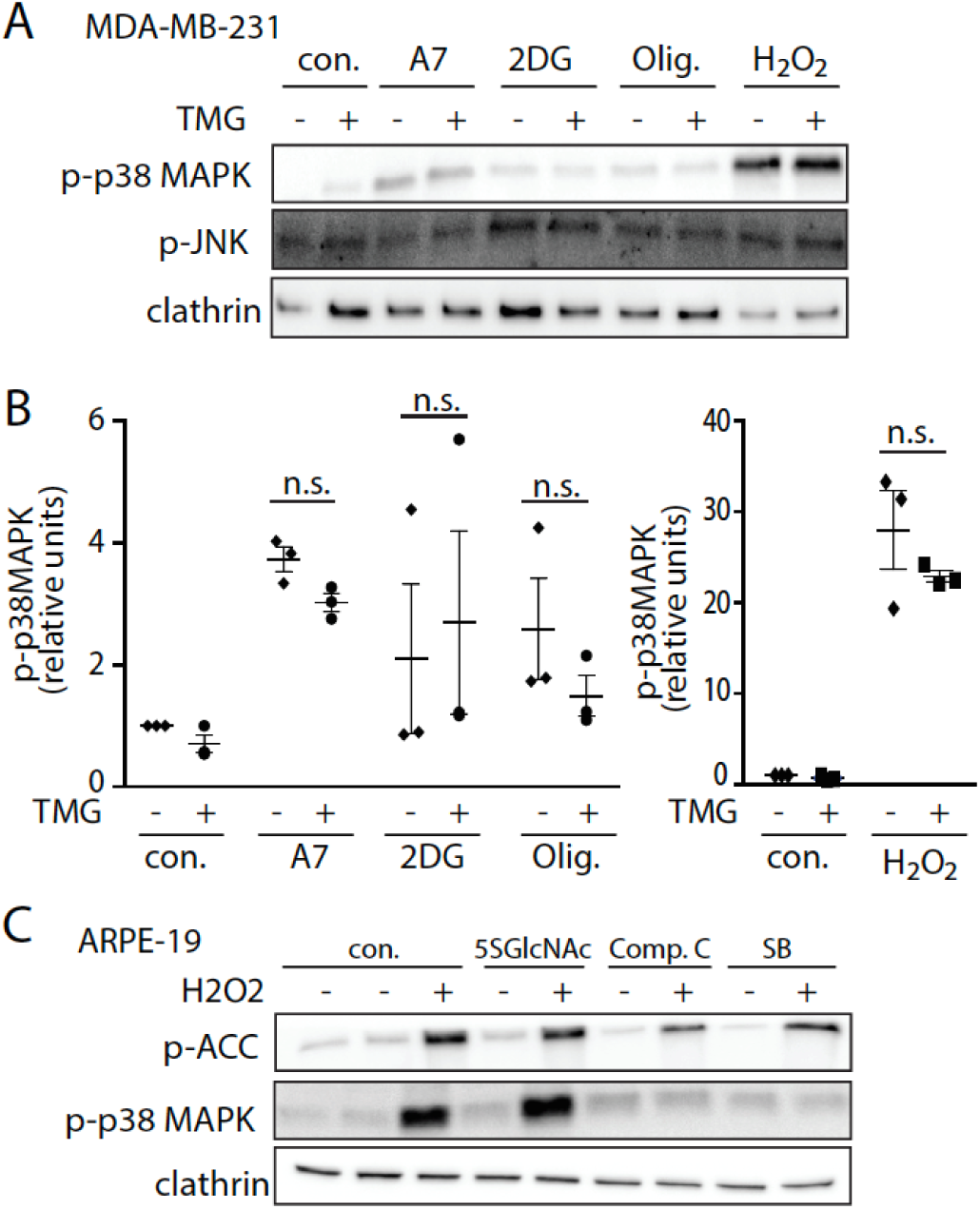
p38 MAPK activation upon treatment with metabolic stress triggers is not impacted by OGT or OGA inhibition. **(A-B**) MDA-MB-231 cells were incubated in low serum for 1 h and then stimulated with either 25 mM 2DG for 20 min, 100 µM A7 for 1 hr, 5 µM oligomycin for 20 min, or 1 mM H_2_O_2_ for 20 min. Representative immunoblots detecting phosphorylated p38 MAPK (p-p38 MAPK), phosphorylated JNK (pJNK) or clathrin (loading control) are shown in (A). Also shown are the means ± S.E. (bars) as well as individual experiment measurements (*n =* 3) for measurements of p-p38 MAPK levels. n.s. (not significant) based on p<0.05 threshold determined by two-way ANOVA with Sidak’s multiple comparisons post-hoc test. (**C**) ARPE-19 cells were incubated in low serum media for 1 h before stimulated with Ac-5SGlcNAc for 4 h, 40 µM Compound C (Comp. C) or 10 µM SB202190 (SB) for 20 min. Shown are representative immunoblots of phosphorylated p38 MAPK (p-p38 MAPK), phosphorylated ACC (pACC) or clathrin (loading controls).

As H_2_O_2_ treatment induced the most potent activation of p38 MAPK, we focused on this treatment for some additional experiments. Consistent with a lack of effect of pre-treatment with the OGA inhibitor Thiamet G, pre-treatment of ARPE-19 cells with the OGT inhibitor 5-thio GlcNAc did not impact H_2_O_2_-stimulated p38 MAPK phosphorylation (**Fig. 5C**). Phosphorylation of p38 MAPK can occur independently of the canonical MKK3/6 signaling mechanism upon exposure of cells to certain cues (Ge et al. 2002), including metabolic stress in which AMPK activation in these conditions leads to p38 MAPK auto-phosphorylation by a TAB1-dependent mechanism (Li et al. 2005; Lanna et al. 2014). H_2_O_2_-stimulated p38 phosphorylation was abolished by treatment with the AMPK inhibitor compound C or the p38 MAPK inhibitor SB202190, suggesting that upon H_2_O_2_ treatment, p38 MAPK activation is dependent on AMPK and p38 MAPK auto-phosphorylation (**Fig. 5C**). That inhibition of either OGT or OGA do not impact p38 MAPK induced by various metabolic stressors (**Fig. 5**) is thus consistent with the lack of effect of OGT or OGA inhibition on AMPK activation (**Fig. 4**), as at least some metabolic stressors (e.g. H_2_O_2_) trigger p38 MAPK phosphorylation in an AMPK-dependent manner.

## DISCUSSION

We find that the ability of a cell to respond to metabolic stressors by increasing global protein O-GlcNAcylation correlates with a lower level of intrinsic ‘basal’ global protein O-GlcNAcylation. As such, cells with elevated levels of O-GlcNAcylation (MDA-MB-231 cells at rest or ARPE-19 cells treated with Thiamet G for 4 hours) exhibited no increase in global protein O-GlcNAcylation upon treatments that trigger metabolic stress signals and AMPK activation. In contrast, these treatments were able to increase global protein O-GlcNAcylation in ARPE-19 cells at rest, and these cells had lower initial global protein O-GlcNAcylation. Together, these results indicate that the extent of initial O-GlcNAcylation may define the extent of further upregulation of global O-GlcNAcylation, such that cells closer to a possible “saturation” of O-GlcNAcylation of target residues on a range of substrates may not be able to further respond to signals that enhance OGT activity, suppress OGA activity and/or enhance UDP-GlcNAc levels.

The increase in global protein O-GlcNAcylation that we observe in ARPE-19 cells upon treatment with A769662 or 2DG for < 1 h is consistent with previous reports that metabolic stress cues and AMPK activation can increase global protein O-GlcNAcylation (Martinez et al. 2017). It is not clear how acute (20 min to 1 h) of metabolic stress can enhance O-GlcNAcylation. AMPK activation enhances glucose uptake in many cell types (Antonescu et al. 2013; Rahmani et al. 2019), and enhanced glucose and glutamine uptake lead to enhanced global protein O-GlcNAcylation (Swamy et al. 2016). Future studies that examine whether the enhancement of global protein O-GlcNAcylation observed in ARPE-19 cells upon treatment with metabolic stressors may be due to enhancement of the uptake of glucose or other nutrients will reveal important insight into this regulation. The signals that may link metabolic stress signals and/or AMPK activation to enhancement of OGT, inhibition of OGA, and/or enhancement of UDP-GlcNAc levels upon nutrient stress, alongside the initial extent of global O-GlcNAcylation prior to nutrient stress, may define the magnitude of the increase in global O-GlcNAcylation in response to metabolic stress.

In contrast to these studies, others have found that global protein O-GlcNAcylation is decreased by AMPK activation, as a result of AMPK-dependent phosphorylation of GFAT1, leading to suppression of the HBP (Eguchi et al. 2009; Zibrova et al. 2017; Gélinas et al. 2018). The relative expression and/or activity of GFAT1 and availability of nutrients such as glucosamine that can bypass the need for GFAT1 in various cell lines may define whether GFAT1 is rate limiting for UDP-GlcNAc production (Fardini et al. 2013; Deen et al. 2016), and thus whether AMPK regulation on GFAT1 can suppress the HBP and global protein O-GlcNAcylation. Some cancer cells exhibit increased levels of GFAT1 expression (Dong et al. 2016; Li et al. 2017) and oncogenic signaling such as that of the epidermal growth factor receptor increases GFAT1 expression (Paterson and Kudlow 1995). Therefore, the ability of AMPK to sufficiently curtail cellular GFAT1 activity may be impaired if GFAT1 expression is substantially elevated in some transformed cells in culture. Furthermore, since cell culture media often has an abundance of nutrients very different to that available to cells in most tissues *in vivo* (Cantor et al. 2017; Voorde et al. 2019), it is conceivable that cells in culture may not have to rely on GFAT1 as the rate limiting step of the HBP in all contexts. Consistent with this interpretation, activation of AMPK with A769662 or phenformin in mouse cardiac tissues resulted in a suppression of global O-GlcNAcylation (Gélinas et al. 2018). This suggests that physiological nutrient levels of O-GlcNAcylation in whole animals may lead to GFAT1 being rate limiting for UDP-GlcNAc production, thus allowing AMPK-dependent phosphorylation and suppression of GFAT1 to reduce UDP-GlcNAc levels and global protein O-GlcNAcylation. Defining the contribution of individual nutrients and/or regulation of metabolic enzymes in establishing GFAT1-dependent control of UDP-GlcNAc production is beyond the scope of the current study but will be important to further establish the parameters that allow AMPK-dependent regulation of global O-GlcNAcylation.

We also find that AMPK activation in ARPE-19 and MDA-MB-231 cells is not dependent on acute (<4 hours) changes in OGT and OGA activity. In contrast, the increase in AMPK activity upon treatment of C_2_C_12_ myotubes with AICAR or by glucose deprivation was blunted by OGA inhibition by Thiamet G or by inhibition of AMPK itself (Bullen et al. 2014). As AMPK can be subject to O-GlcNAcylation on both α and all three γ subunits, Bullen et al. suggest that dynamic cycles of AMPK modification with O-GlcNAc and hydrolysis of O-GlcNAc were required for AMPK activation (Bullen et al. 2014). In contrast, in ARPE-19 cells and MDA-MB-231 cells, we did not observe any effect of inhibition of OGT or OGA on AMPK activation, measured by phosphorylation of the AMPK substrate ACC, in response to treatment with A769662, 2DG, or H_2_O_2_. This establishes that O-GlcNAcylation is not strictly required for the regulation of AMPK activation upon exposure to metabolic stress triggers, and that instead O-GlcNAcylation may establish control over AMPK only in specific contexts.

What conditions may allow O-GlcNAcylation to effect control over AMPK? In proliferating cells, OGT was observed to have a strong nuclear localization, while OGT also accumulated in the cytosol in differentiated cells such as C_2_C_12_ myotubes (Bullen et al. 2014). Thus, it is possible that localization of OGT and/or AMPK to distinct compartments (such as strong nuclear localization of OGT in proliferating cells) prevent regulation of AMPK by O-GlcNAcylation. Indeed, ARPE-19 and MDA-MB-231 cells are rapidly proliferating, suggesting that they may exhibit significant nuclear localization of OGT. However, our results indicate that O-GlcNAcylation is not an essential contributor to AMPK activity in all contexts but instead may modulate AMPK activation specifically in some cellular contexts, such as in differentiated cells like C_2_C_12_ myotubes (Bullen et al. 2014).

In addition, as AMPK is activated by allosteric mechanisms upon AMP/ADP binding, as well as binding of AMPK to a lysosomal complex containing the v-ATPase, aldolase and LKB1 in absence of the glycolytic intermediate fructose-1,6-bisphosphate (Zhang et al. 2017). This suggests that AMPK activation mechanisms may differ in response to different metabolic signals, leading to distinct regulation of AMPK by reductions of ATP/AMP versus reduced glycolysis and fructose-1,6-bisphosphate. However, we note that we observe a lack of regulation of AMPK by inhibition of OGT and OGA in response to diverse mechanisms of activation of AMPK: A769662 (an allosteric activator that binds to the β1-subunit of AMPK,(Sanders et al. 2007; Xiao et al. 2013; Calabrese et al. 2014)), 2DG (which reduces glycolytic flux, (Pajak et al. 2020)), and H_2_O_2_ (which induces reactive oxygen species stress that can impact AMPK directly, (Hinchy et al. 2018), or indirectly as a disruption of mitochondrial metabolism, (Hinchy et al. 2018). As such, our results suggest that it is unlikely that distinct cues leading to AMPK activation in a specific cell type are differentiated by distinct regulation by O-GlcNAcylation. Nonetheless, it remains possible that O-GlcNAcylation selectively controls upstream activators of AMPK such as LKB1 or CAMKII (Dias et al. 2012), such that specific cells and/or contexts that rely on specific O-GlcNAc modified protein(s) for AMPK phosphorylation or dephosphorylation would exhibit regulation of AMPK activation indirectly.

Our results indicate that AMPK activation enhances global O-GlcNAcylation in ARPE-19 cells but not in MDA-MB-231 cells. In addition to control of global O-GlcNAcylation levels, AMPK may regulate the substrate specificity of OGT, perhaps due to AMPK-dependent phosphorylation of OGT on Thr444 (Bullen et al. 2014). We did not examine the O-GlcNAcylation of specific substrates here, focusing instead on measurement of global O-GlcNAcylation, a common measurement of OGT activity (Zachara et al. 2004; Swamy et al. 2016; Zhu et al. 2018; Gélinas et al. 2018). In addition to control of OGT by direct AMPK-dependent phosphorylation that regulates OGT substrate specificity, the substrate specificity of OGT is also controlled by p38 MAPK via direct binding of p38 MAPK to OGT in a p38 MAPK activity-dependent manner (Cheung and Hart 2008). Since p38 MAPK is activated by a variety of metabolic stress signals, indeed in many instances in an AMPK-dependent manner (Li et al. 2005; Lanna et al. 2014) as we have shown here, this further supports the notion (Bullen et al. 2014) that metabolic stress signals may also alter AMPK substrate specificity. Nonetheless, our work establishes that in addition to control of OGT substrate specificity, metabolic stress signals also enhances global protein O-GlcNAcylation in a manner that correlates with sub-saturating levels of initial global O-GlcNAcylation.

We have focused our studies on acute (<4 hours) treatments with inhibitors of OGT or OGA and treatment with metabolic stress triggers. Our experiments suggest that AMPK activation and global protein O-GlcNAcylation can be orthogonal; however, the interplay between these two signals may occur at longer time-points. At longer time points of treatment with metabolic stress signals and AMPK activation and/or treatment with inhibitors of OGT/OGA, additional layers of regulation such as transcriptional control of OGT and OGA (Kim et al. 2019) may reveal additional possible cross-talk of AMPK and O-GlcNAcylation signaling (Laarse et al. 2018)

The activation of AMPK and regulation of global and/or substrate-specific protein O-GlcNAcylation have important contributions in allowing cellular adaptation to specific metabolic conditions in various physiological and pathophysiological contexts, as has been recently reviewed (Rahmani et al. 2019). For example, these signals are important to allow cancer cells to modulate their metabolism to adapt to stressful tumor microenvironments (Fardini et al. 2013; Martinez et al. 2017), such as through control of lipid metabolism observed in MDA-MB-231 breast cancer cells (Sodi et al. 2018). Since AMPK and O-GlcNAcylation can exhibit orthogonal activation and signaling (this study), this suggests that understanding how each of these signals may sense distinct physiological metabolic cues and/or target distinct adaptive outcomes will be important in understanding diseases such as cancer.

Cells harbour many signaling pathways to control the activity of proteins in various physiological time scales under different contexts to allow adaptation of cell physiology to meet a range of conditions. There are a number of signals that respond to changing nutrient conditions to promote cellular adaptation. Whether these metabolic stress signaling pathways exhibit obligatory cross-talk or whether they can operate orthogonally under some circumstances to respond to nutrient limitation or sufficiency is only beginning to be understood. Our results thus suggest that AMPK and O-GlcNAcylation can be orthogonal signals that are modulated by nutrient availability and/or metabolic stress triggers and exhibit reciprocal regulation only under specific cellular contexts. This highlights the complexity of metabolic parameters that must be sensed at the cellular level and the diversity of metabolic adaptations for optimal cellular and systemic survival.

## AUTHOR STATEMENTS

### Acknowledgements

We are grateful to Dr. David Vocadlo at Simon Fraser University (Canada) for the kind gift of 5-thio GlcNAc (peracetylated 2-acetamido-2-deoxy-5-thio-D-glucopyranose, Ac-5SGlcNAc).

### Author Contributions

S.R.: conceptualization, investigation, formal analysis, writing; M.H.: investigation, formal analysis, W.W.W.: conceptualization, supervision, funding acquisition, writing, C.N.A.: conceptualization, supervision, resources, project administration, funding acquisition, formal analysis, writing.

