## Supplemental Materials for "Differences in intrinsic cellular O-GlcNAcylation impact response to metabolic stress"

**Supplemental Figures:**

**Figure S1:** The O-GlcNAc antibody specifically recognizes O-GlcNAc modified nucleocytoplasmic proteins

**Table S1:** List of key reagents and resources used in this study

**Materials:** siRNA gene silencing of ARPE-19 cells

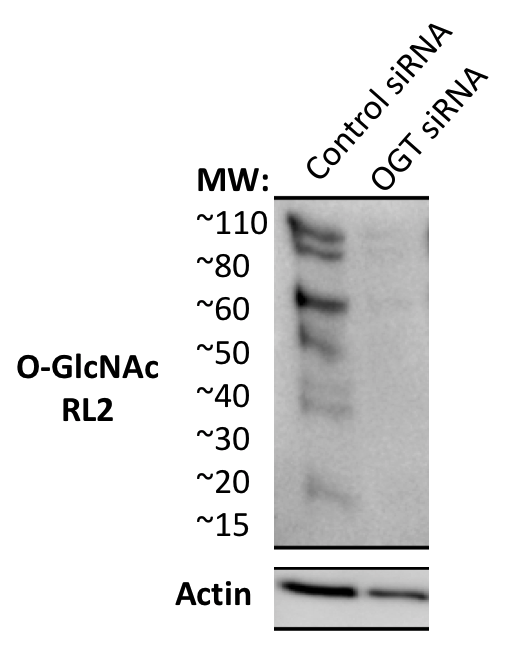

**Figure S1. The O-GlcNAc antibody specifically recognizes O-GlcNAc modified nucleocytoplasmic proteins.**

O-GlcNAc transferase (OGT), which modifies nucleocytoplasmic proteins with O-GlcNAc, was silenced in ARPE-19 cells using siRNA gene silencing. Whole-cell lysates were prepared and probed with RL2 O-GlcNAc antibody (Abcam, ab2739) to detect proteins modified with O-GlcNAc residue. Shown is the representative blot indicating that this antibody used in our study is specific to O-GlcNAc modified proteins at all molecular weights (MW).

**Table S1. List of key reagents and resources used in this study.**

| **Reagent or resource** | **Source** | **Product code** |
| --- | --- | --- |
| **Antibodies** | | |
| p-ACC S79 (D7D11) | Cell Signaling Technology | 11818S |
| p-p38 MAPK T180/Y182 (D3F9) | Cell Signaling Technology | 4511S |
| Clathrin (D3C6) XP | Cell Signaling Technology | 4796S |
| Pan Actin (D18C11) | Cell Signaling Technology | 8456S |
| Anti-O-Linked N-Acetylglucosamine antibody [RL2] | Abcam | ab2739 |
| p-SAPK/JNK T183/Y185 | Cell Signaling Technology | 9251S |
| Anti-mouse HRP | Cell Signaling Technology | 7076 |
| Anti-rabbit HRP | Cell Signaling Technology | 7074 |
| **Chemicals** | | |
| A769662 (A7) | Abcam | ab120335 |
| Oligomycin (OG) | Cell Signaling Technology | 9996 |
| 2-deoxy-D-Glucose (2DG) | Sigma Aldrich, Oakville, ON | 1002939980 (D6134) |
| Hydrogen peroxide | Bio Basic | HC4060 |
| Thiamet G (TMG) | Sigma Aldrich | SML0244  (Batch 22957) |
| Compound C | Abcam | Ab120843 |
| SB202190 | Sigma Aldrich | S7067 |
| DMSO | BioShop, Burlington, ON, Canada | 276855 |
| 5-thio-GlcNAc (5-S-GlcNAc) | Simon Fraser University, Burnaby, BC, Canada | Reference: (Gloster, Zandberg, Heinonen, Shen, & Vocadlo, 2011) |
| **Experimental Models: Cell Lines** | | |
| ARPE-19 | ATCC | RRID: CVCL_0145 |
| MDA-MB-231 | ATCC | RRID: CVCL_0062 |

**Methods for Fig S1:**

**siRNA gene silencing of ARPE-19 cells:** ARPE-19 cells were subject to siRNA gene silencing for O-GlcNAc transferase and non-targeting control. Custom siRNAs were synthesized to target specific transcripts with sequences as follows: OGT: GAA GAA AGU UCG UGG CAA A

(sense strand) and non-targeting CGU ACU GCU UGC GAU ACG GUU (sense strand). siRNA gene silencing (Fig. S1) was performed using Lipofectamine RNAiMAX (Life Technologies, Carlsbad, CA), as per the manufacturer’s instructions and as previously described (Garay et al., 2015). Each siRNA construct was transfected at 220 pmol/l precomplexed and incubated in in the transfection reagent in Opti-MEM (Gibco) for 4 hrs. After the 4 hr incubation, cells were washed and replaced in regular 10% FBS DMEM/F12 growth medium. siRNA transfections were performed twice (72 hr and 48 hr) before whole-cell lysate preparation.
